# PureCLIP: capturing target-specific protein-RNA interaction footprints from single-nucleotide CLIP-seq data

**DOI:** 10.1101/146704

**Authors:** Sabrina Krakau, Hugues Richard, Annalisa Marsico

## Abstract

iCLIP and eCLIP techniques facilitate the detection of protein-RNA interaction sites at high resolution, based on diagnostic events at crosslink sites. However, previous methods do not explicitly model the specifics of iCLIP and eCLIP truncation patterns and possible biases. We developed PureCLIP, a hidden Markov model based approach, which simultaneously performs peak calling and individual crosslink site detection. It explicitly incorporates RNA abundances and, for the first time, non-specific sequence biases. On both simulated and real data, PureCLIP is more accurate in calling crosslink sites than other state-of-the-art methods and has a higher agreement across replicates. Link: **https://github.com/skrakau/PureCLIP**.

## Background

Interactions between RNAs and RNA binding proteins (RBPs) play essential roles in both transcriptional and post-transcriptional gene regulation. RBPs bind on several sites of both coding and non-coding RNAs with directed but somewhat fuzzy binding affinity for both RNA sequence and structure. In order to fully understand regulatory processes mediated by RBPs it is crucial to accurately determine the full landscape of interactions for a protein of interest. State-of-the-art technologies using crosslinking and immunoprecipitation combined with high-throughput sequencing (CLIP-seq) allow a genome-wide binding site detection with high resolution. The most commonly used protocols in this field are HITS-CLIP [6], photoactivatable-ribonucleoside-enhanced CLIP (PAR-CLIP) [13] and since 2010 individual-nucleotide CLIP (iCLIP) [15]. All methods use UV light, which causes the formation of crosslinks at protein-RNA interaction sites. These crosslinks subsequently increase the probability for base transitions, deletions and truncations during the reverse transcription. Such *diagnostic events* can be used to localize the crosslink position. However, due to the ligation of an adapter at the 5' end of the RNA fragments, HITS-CLIP and PAR-CLIP methods only capture cDNAs which are entirely read by the reverse transcriptase, i.e. not truncated. The fraction of truncated and thus lost fragments is typically over 80% [27].

iCLIP-seq uses a cleavable adapter in combination with an additional circularization step, which allows all cDNA fragments to be amplified and sequenced. As a consequence, valuable information about the exact crosslink site can be retained from truncated cDNAs, or more precisely from the read start sites they cause. Recently, various improvements to the protocol were proposed to alleviate previous limitations [12,24]. Another protocol called eCLIP was published in 2016 [30]. Similarly to iCLIP, it provides single-nucleotide resolution by capturing truncated cDNAs but, due to the optimization of several steps, improves the specificity of called binding sites. To date, eCLIP datasets for more than 120 different proteins have been published as part of the ENCODE consortium [3,25]. While previous CLIP-seq experiments often had matched IgG control experiments, which suffer from sparsity and high amplification rates [30], the eCLIP-seq protocol is designed to generate a size-matched input control. This input control is sampled prior to the immunoprecipitation and thus contains the signal of non-specific background.

In order to infer target-specific RBP binding regions from iCLIP/eCLIP data, it is crucial to account for different sources of biases, such as transcript abundances, crosslinking sequence preferences [27] and mappability. The crosslinking sequence bias can also be observed within the eCLIP input data, since it “represent[s] RNAs crosslinked to many different RBPs and should reflect the sequence preferences at crosslink sites that are common to a mixture of RBPs” [12]. Haberman et al. showed that certain polypyrimidine rich k-mers, which they call crosslink-associated (CL) motifs are enriched at read start sites in both input and target eCLIP data compared to upstream regions [12].

Besides background noise such as signal coming from sticky RNA fragments or non-specific crosslink events within CL-motifs, the binding of background proteins constitute a major challenge for the analysis of CLIP-seq data. A recent study analysing previously published PAR-CLIP datasets showed that if no control dataset is used for correction, up to 45% of the called binding sites overlap with background binding sites [9]. Background binding regions that are common to several CLIP-seq datasets have been systematically identified [21] and can be used for validation of called binding sites. These findings demonstrate the importance of control experiments, such as input experiments, to reduce the number of false positives at such regions.

Several tools have been developed for the computational analysis of HITS-CLIP and PAR-CLIP data [4, 23,29], but very few tools have been developed which are tailored for the specific analysis of iCLIP/eCLIP data. In addition, previous methods for CLIP-seq data analysis do not fully take into account possible sources of bias, such as transcript abundances and non-specific CL-motifs, which heavily affect iCLIP and eCLIP data [12,28], thereby returning a high number of false calls. The tool Piranha [29] performs strand-specific peak-calling without explicitly normalizing for non-specific background signal. It models the underlying bin-wise read count distribution to compute a genome-wide significance threshold above which peaks are called. CLIPper [30] is also a strand-specific peak-calling method designed by members of the ENCODE consortium for the analysis of the published eCLIP datasets. It incorporates annotations from the reference genome and computes significance thresholds on a gene-by-gene basis. Both tools, Piranha and CLIPper are peak calling methods which do not detect individual crosslink sites. Their limitation is that they potentially miss low affinity binding regions with a clear iCLIP truncation pattern due to the arbitrary setting of a threshold on the number of reads. In addition, they are sensitive to call peaks caused, for example, by artefacts within high abundant RNAs. The method CITS on the other hand aims to call individual crosslink sites from iCLIP-seq data [31]. It clusters reads based on their start sites and uses a statistical test to detect sites within such clusters containing a significant fraction of read starts. A drawback of this method is that it does not explicitly model the relation between read start counts and the read coverage generated by pulled-down iCLIP fragments. As a result, it might be also sensitive to artefacts within highly abundant RNAs. In contrast, PIPE-CLIP [2] is an online pipeline for the analysis of HITS-CLIP, PAR-CLIP and iCLIP data designed to separately call peaks and crosslink sites, which are subsequently merged. Although constituting a powerful idea, one drawback of this method is that it is not designed to include control experiments in the analysis. In addition, being designed to be an online method, its application for transcriptome-wide analysis is not practically feasible. As described above, both CLIP-seq peak calling methods and individual crosslink site detection methods come with advantages and disadvantages, but currently no method exists which addresses peak-calling and individual crosslink site detection simultaneously, while correcting for possible biases.

We have developed PureCLIP, a method to capture target-specific protein-RNA interaction footprints from iCLIP/eCLIP-seq data. PureCLIP calls individual crosslink sites considering both regions enriched in protein-bound fragments and iCLIP/eCLIP specific truncation patterns. Our method uses a non-homogeneous Hidden Markov model to incorporate additional factors into the model, such as non-specific background signal from input experiments and CL-motifs in order to reduce the number of false positives. We have exhaustively validated the superiority of PureCLIP over several existing methods in various settings. First, we have designed a realistic iCLIP/eCLIP simulation setup and demonstrated that, over a wide range of simulation parameters, PureCLIP is up to 7-15% more precise than other methods in detecting target-specific crosslink sites. Second, in the lack of an experimental gold standard, we have selected 4 datasets of published iCLIP/eCLIP data for evaluation where the RBP motif or the predominant binding region of the RBP is known. We consistently observed that PureCLIP is better than other methods in determining *bona fide* binding site locations. In particular, the incorporation of covariates, such as input signal and CL-motifs, increases the precision of PureCLIP up to 8-10% compared to previous methods. Third, the replicate agreement of target-specific crosslink sites called by PureCLIP is up to 8-20% higher than other methods, indicating that PureCLIP is highly specific in crosslink site detection.

## Results

### Overview of the approach

PureCLIP aims to detect individual crosslink sites originating from interactions between RNAs and the protein targeted by the experiment. In order to accomplish this, we address two objectives: (1) detect regions enriched in mapped reads caused by pulled-down RNA fragments and (2) detect crosslink sites where a significant fraction of read starts accumulate at the same position, originating from truncated cDNAs (Fig. 1).

In the following we give an overview on how we derive this information, assuming that the input data are iCLIP/eCLIP-seq reads which have been mapped to either a genome or a transcriptome and PCR duplicates have been removed. The output of PureCLIP consists of individual crosslink sites associated with a score. Since multiple crosslink sites can occur within one binding region, the crosslink sites are optionally merged.

### Hidden Markov model

CLIP-seq data features a spatial dependency between neighbouring positions. In order to infer crosslink sites from the observed data we look at it as a segmentation problem and address this using a hidden Markov model (HMM). The HMM has a single-nucleotide resolution and each position can be categorized either as 'non-enriched' or 'enriched', indicating whether the position is enriched or not in protein bound fragments. In addition, each position can also be categorized as 'non-crosslink' or 'crosslink', indicating whether it represents a crosslink site or not. This combination results in four hidden states: (1) 'non-enriched & non-crosslink', (2) 'non-enriched & crosslink', (3) 'enriched & non-crosslink' and (4) 'enriched & crosslink' (Fig. 1). State (2) corresponds to non-specific crosslink sites and it is included in the model for mathematical completeness. We are interested in all sites with a hidden state (4), i.e. sites that are enriched in pulled down RNA fragments and show the truncation pattern (Fig. 1a).

**Figure 1:**
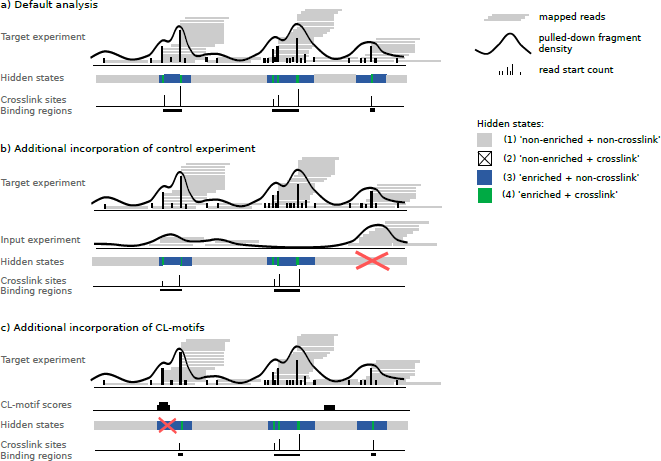
**Overview of PureCLIPs approach. a)** PureCLIP starts with mapped reads from a target iCLIP/eCLIP experiment and derives two signals: the pulled-down fragment density and individual read start counts. Based on these two observed signals it infers for each position the most likely hidden state. The goal is to identify all sites with an 'enriched & crosslink' state. Individual crosslink sites can then be merged to binding regions. **b)** Additionally, information from input control experiments can be incorporated. The fragment density is used to correct for non-specific background signal, which reduces the number of false calls. **c)** Furthermore, PureCLIP can incorporate information about CL-motifs, in order to reduce false calls caused by non-specific crosslinks.

In order to detect 'enriched & crosslinked' sites, PureCLIP uses two signals derived from the mapped reads: (1) the *pulled-down fragment density,* which is a smoothed signal derived from the read start counts and holds information about the enrichment within the current region; (2) and the read start counts themselves, which hold information about potential truncation events (Fig. 1). One type of distribution is used to model the pulled-down fragment density, with one set of parameters for the 'non-enriched' state and one for the 'enriched' state, assuming that the 'enriched' state is more likely to cause high fragment density values than the 'non-enriched' state. Similarly, read start counts are modelled under the assumption that the 'crosslink' state is more likely to generate a higher fraction of reads starting at one position than the 'non-crosslink' state. In order to account for differently covered regions, the parameters of the read start count distributions at individual positions depend on the pulled-down fragment density.

The fragment density distributions and the read start count distributions are combined to obtain the emission probabilities of each of the four hidden states. For each position we can then address the question: which of the four hidden states did most likely cause the observed data?

### Incorporation of additional factors into the PureCLIP model

The observed signals can be biased by a number of different factors, such as transcript abundance or crosslinking sequence preferences. An important feature of PureCLIP is the incorporation, into the HMM framework, of position-wise external data to correct for such biases. We do this by using generalized linear models (GLMs) and model this separately for different types of covariates.

We expect regions within highly abundant RNAs to show more read start counts than regions within less abundant RNAs. This holds for both target binding regions and for regions with non-specific background noise. To normalize for this, information from input control experiments can be included to influence the emission probability distributions of the 'non-enriched' and 'enriched' states. With this, we aim to reduce the number of false positives for highly abundant RNAs (see Fig. 1b) while increasing the sensitivity for less abundant RNAs.

Furthermore, we expect a higher number of read start counts, for example, at positions within CL-motifs. Thus, in order to correct for the crosslinking sequence bias, information about CL-motifs can be incorporated (see Fig. 1c) to influence the 'non-crosslink' and 'crosslink' emission distributions.

## Evaluation of PureCLIPs performance in comparison to previous strategies

Evaluating a method's performance in analysing CLIP-seq data is not trivial, since no gold-standard of binding regions or crosslink sites exists. We addressed this task by (1) assessing the precision and recall of PureCLIP in basic mode, i.e. without additional covariates, in calling individual crosslink sites on simulated data. (2) We then used real iCLIP and eCLIP datasets of proteins with known binding characteristics, such as known sequence motifs or known predominant binding regions. We assess the ratio of sites called by each method which falls within these motifs or inside those binding regions. Called crosslink sites within such regions are defined as true positives. Here we applied PureCLIP in four different settings: in basic mode, incorporating input signal, incorporating CL-motifs and incorporating both input signal and CL-motifs simultaneously. Although extremely valuable, this evaluation approach is limited by the fact that it is unknown how far the protein of interest can also bind to alternative motifs or outside the defined *bona fide* binding regions. For this reason, (3) we also assessed the agreement of called crosslink sites between eCLIP replicates.

We compared PureCLIP against a range of previous strategies, most importantly CITS [31], which similarly to PureCLIP can call individual crosslink sites rather than broader peak regions. Additionally, since to this date, no other tool exists that addresses both peak calling and crosslink site detection simultaneously for truncation based CLIP-seq data, we combine the peak-calling methods Piranha [29] and CLIPper [17] with CITS. More precisely, we use the intersection of the called peaks and the CITS crosslink sites. While this intersection depends on the selected p-value thresholds for both methods, the resulting sites are scored in two different ways, using either the score from the peak-calling method (referred to with the term Piranha^sc^ or CLIPper^sc^) or from CITS (referred with CITS^sc^) (for details see Materials and Methods). With this we aim to cover the range of currently available strategies for detecting protein-RNA interactions at single-nucleotide resolution. To ensure a comparative assessment which is as impartial as possible, we also compared PureCLIP also with combination based on different p-value thresholds and found that it does not affect the results (see Additional file 1, Fig. S4).

Additionally, we applied the simplest possible approach, namely calling all sites with a read start count above a certain threshold. This gives us an understanding of how different methods perform in different scenarios compared to this naive approach. In the following we refer to this as the *simple threshold* method.

## PureCLIP outperforms previous strategies on simulated iCLIP/eCLIP-seq data

Since the only available CLIP-seq simulator [14] is limited to PAR-CLIP and HITS-CLIP data, we implemented our own simulation workflow to mimic the experimental steps of iCLIP and eCLIP protocols. Starting from real RNA-seq data and known binding regions of a certain protein, our simulation aims to reproduce the characteristics of iCLIP/eCLIP data as accurately as possible. To simulate target signal, our workflow uses aligned RNA-seq data, 'pulls-down' a certain fraction of the fragments that cover a known binding region, and then applies truncations according to a given rate (for more details see Methods). Furthermore, nonspecific binding of background proteins is simulated -using published common background regions, and random noise from RNA-seq data is added.

To evaluate PureCLIP's performance under different conditions, we produced three different datasets. For these we used varying 'pull-down' rates for the target signal, i.e. either 100% or 50% of the RNA fragments that overlap a target binding region are selected and further modified where required. Reducing the pulldown rate enables us to get an idea how the different methods perform for proteins with overall lower binding affinities. Additionally, we simulated non-specific background binding for two of the datasets (see Fig. 2).

For the evaluation, we define a called crosslink site as a true positive if a target crosslink site was simulated at the same position. The precision of a method is calculated as the fraction of true positives among the called crosslink sites. We first investigated the precision versus the number of true positive crosslink sites. The results in Figure 2 (top) demonstrate that PureCLIP reaches a higher precision in detecting individual crosslink sites than previous strategies for all simulation settings. In particular for the top ranking sites, it has a far better precision compared to other methods, while being comparable to 'CITS^sc^ + CLIPper' for more sensitive settings. However, it is worth mentioning here, that sensitive settings which are characterized by a precision below 50% are generally not of interest.

Furthermore, we investigated whether the crosslink sites called by PureCLIP could be used to recover target binding regions (i.e. known binding regions in which crosslink sites were simulated) or if they cluster within a few regions with high fragment density. A target binding region is counted as a true positive, if it could be recovered with at least one called crosslink site. The precision is defined as the percentage of called crosslink sites within target binding regions. For all simulation settings the results show that PureCLIP recovers binding regions with higher precision compared to previous strategies (Fig. 2, bottom).

Note that our method depends on the bandwidth used for the smoothing of the read start counts. The optimal bandwidth depends on the coverage and the given cDNA length distribution, e.g. the longer the cDNAs the larger the optimal bandwidth. For this evaluation we used a bandwidth of 50 bp, which is also used for the real data analysis. The results shown in Fig. S5 in Additional file 1 demonstrate that PureCLIP reaches a higher precision robustly for a range of different bandwidth parameters, compared to previous strategies.

## PureCLIP detects *bona fide* binding regions with higher precision compared to previous strategies

We used publicly available eCLIP (PUM2, RBFOX2, U2AF2) [30] and iCLIP (U2AF2) [32] datasets to measure the performance of the different strategies in calling crosslink sites within *bona fide* binding regions. For PUM2 and RBFOX2 these binding regions were defined by their known sequence motifs (see Additional file 1, Fig. S1), while for U2AF2 we make use of its known predominant binding region ~ 11 nt upstream of 3'splice sites [32]. Here, a sequence motif based definition of the binding region is not applicable, since U2AF2 binds to poly(U) tracts, which coincide with non-specific CL-motifs.

For the PUM2 data, all strategies revealed an accumulation of called crosslink sites at the 5' end of PUM2 motif occurrences and another slightly weaker accumulation towards the 3' end (Fig. 3a, left panel). For RBFOX2 eCLIP data we observe an accumulation of called crosslinks at the two guanines within the motif (Fig. 3b, left panel). These crosslinking patterns are in agreement with previous studies [31] [30] and, since crosslinks do not preferentially occur at guanines, are most likely caused by target-specific protein-RNA interactions.

### PureCLIPs performance without incorporating external data as covariates

We first investigated the precision of PureCLIP in basic mode, i.e. without the incorporation of any covariates, where calls are considered true positives if they fall within the motif area or upstream of 3' splice sites. We observed that PureCLIP outperforms all other methods even without covariates in three out of four datasets, as shown in Figure 3 (right panel). Interestingly, when applying strategies that merge results from peak-calling tools and CITS, using the peak-calling scores for ranking ('Piranha^sc^ + CITS', 'CLIPper^sc^ + CITS'), we always get a lower precision than when using the CITS crosslink site detection score for ranking ('Piranha + CITS^sc^', 'CLIPper + CITS^sc^').

### Incorporation of input control data improves crosslink site detection

We expect the observed smoothed read start counts to be biased by different factors, among others by RNA transcript abundances. The published eCLIP datasets come with input control experiments [30], which provide information about the non-specific background signal, i.e. RNA fragments crosslinked to background proteins. We observe significant correlations between the fragment density of the eCLIP target dataset and the input dataset with Pearson correlation coefficients ranging from 0.36 to 0.42 (p-values < 2.2e-16) (see Additional file 1, Fig. S3a). Therefore, the incorporation of input signal into the PureCLIP framework gives us the possibility to indirectly normalize for transcript abundances, crosslinking preferences and other local biases.

In detail, the PureCLIP framework uses the eCLIP input signal to model the emission probabilities of the 'non-enriched' and 'enriched' states for the observed data, i.e. the pulled-down fragment density. This means that instead of using one global emission probability distribution for the 'non-enriched' or 'enriched' states, the position-wise input signal is used to model the expected mean parameter of each of the two emission probability distributions (see Additional file 1, Fig. S3b). With this we aim to reduce the number of false positives, for example, within highly abundant RNAs while increasing the sensitivity within lowly abundant RNAs. The evaluation based on *bona fide* binding regions from real data shows that incorporating the input signal improves the precision of PureCLIP for all eCLIP datasets and over all sensitivity thresholds (Fig. 3 a-c, right panel). In particular for the top ranking sites, it greatly improves the precision by reducing the number of false positives in regions showing high non-specific background signal.

### Incorporation of CL-motifs greatly improves crosslink site detection

Another major bias within CLIP-seq data is caused by crosslinking sequence preferences, which influence the individual read start counts. This also gives rise to non-specific crosslink events at sites with no direct interaction between the target protein and the RNA. Since our method is designed to detect crosslinking patterns, it also detects a certain fraction of non-target crosslink sites. For PUM2 and RBFOX2, both having known sequence binding motifs distinct from reported CL-motifs [12], we observed that 33% and 37% of the top 1000 sites called by the basic version of PureCLIP overlap with regions harbouring a CL-motif.

**Figure 2:**
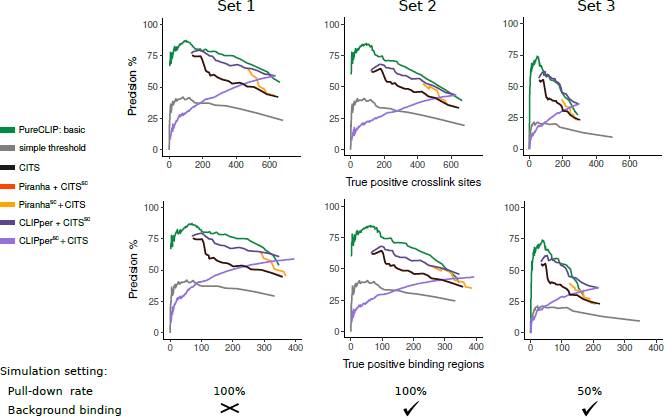
Accuracy on simulated iCLIP-seq data. Precision and number of true positive crosslink sites (top) and of binding regions (i.g. regions with at least one crosslink site, bottom) on three different simulation settings. Characteristics of each simulation are reported at the bottom. The leftmost point of each curve corresponds to the highest precision level that the corresponding method can report. The curves of 'Piranha + CITS^sc^' overlap with CITS.

In order to reduce the number of such potential false positives, we incorporate information about CL-motifs into our model. This can be particularly helpful to filter out non-specific crosslink sites when the protein of interest preferentially binds sequences that are clearly distinct from CL-motifs. For this purpose, CL-motifs have to be learned first and we do this directly from the data: (1) we call crosslink sites in the eCLIP input data, (2) we then learn CL-motifs on these sites using DREME [1] and (3) we apply FIMO [11] to compute the occurrences of those motifs and their scores within the reference genome or transcriptome. These position-wise scores are then incorporated into the HMM framework of PureCLIP to model the emission probabilities of the 'non-crosslink' and 'crosslink' state for the observed data, i.e. the read start counts. This enables a correction for the crosslinking sequence bias at CL-motif positions. As an example, the four most enriched CL-motifs from the analysis of PUM2 eCLIP input data are shown in Fig. 4.

The results demonstrate that for PUM2 (Fig. 3b) and in particularly for RBFOX2 eCLIP data (Fig. 3a) the incorporation of CL-motif scores greatly improves the precision in calling crosslink sites within *bona fide* binding regions. Interestingly, the simultaneous incorporation of the input signal and CL-motif scores improves the precision of PureCLIP even further (Fig. 3a, b). Moreover, we can see that for the protein U2AF2, whose sequence motif coincides with CL-motifs, the performance of PureCLIP stays robust and is not impaired by the incorporation of CL-motif scores. Altogether, we could see that when incorporating CL-motifs PureCLIP consistently performs better than previous strategies in positioning called sites either at the known binding motif or ~ 11 nt upstream of 3' splice sites for U2AF2 (Fig.3a-d).

**Figure 4:**
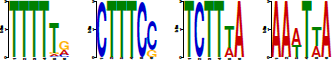
CL-motif analysis of PUM2 eCLIP input data. Logo representation of the four top scoring motifs among the first 5000 PureCLIP crosslink sites called on the input dataset. Motifs were detected with DREME and a 10 bp window around the crosslink sites. As previously reported [12], polypyrimidine rich motifs are overrepresented.

## PureCLIP has a higher agreement of called sites between eCLIP replicates compared to previous strategies

Besides using known binding regions for evaluation, we aimed to assess the performance of the different methods independently of that information, since in the end the exact binding regions and crosslink sites remain unknown. For this reason we explored the agreement of called crosslink sites between eCLIP replicates, assuming that target-specific binding events are more likely to be observed in both replicates than nonspecific noise. We applied all methods to the individual eCLIP replicate datasets and measured for each sensitivity threshold how many of the *x* called crosslink sites in replicate 1 overlap with the top *x* ranking crosslink sites in replicate 2.

**Figure 3:**
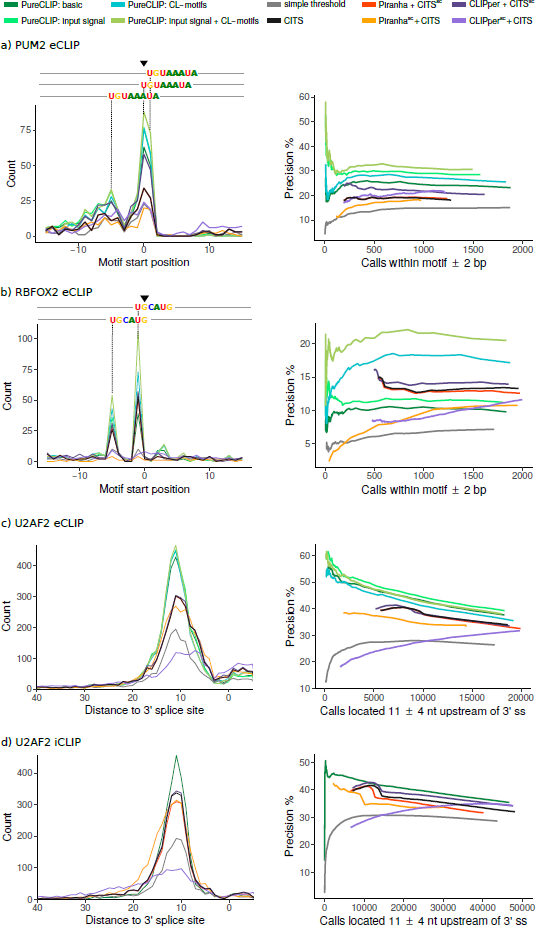
Accuracy in detecting *bona fide* binding regions. **Left panel: a)** distribution of the closest PUM2 motif start positions relative to the crosslink site for the top 1000 sites called by each method. Position 0 represents the crosslink site. **b)** Same as a), but for RBFOX2 motif start positions. **c) - d)** Distribution of the distances of the top 5000 sites called by each method with respect to 3' splice sites. **Right panel:** Precision of the called sites for all methods at different sensitivity settings (not only for the top sites). The leftmost point of each curve corresponds to the highest precision level that the corresponding strategy can report.

We found that besides target-specific binding events also other factors contribute to this replicate agreement, such as the binding of highly abundant background proteins at highly abundant RNAs or the crosslinking sequence bias (see Additional file 1, Section 7). To ensure that the computed replicate agreement represents an unbiased measure of the method's precision, we only consider sites that are enriched over the input and located outside of regions known be prone to background binding (published in [21]). In the following, we refer to this as the *bias corrected* replicate agreement (see Methods for details). To further prevent contribution from common non-specific crosslinks, for PUM2 and RBFOX2 we only counted sites which are not located within CL-motif occurrences. The U2AF2 iCLIP data is excluded from this evaluation, since no input control experiment is available and thus the bias corrected replicate agreement can not be computed.

Our evaluations show that PureCLIP has a higher bias corrected replicate agreement for the top ranking sites compared to previous strategies, in all four PureCLIP settings and over all three eCLIP datasets, as depicted in Figure 5. Furthermore, the performance of PureCLIP in basic mode is at least comparable to other methods, while PureCLIP incorporating input signal and CL-motifs strictly outperforms all other methods. While the individual use of these covariates already improves the agreement, the best results are obtained when both of them are incorporated simultaneously.

Notably, other strategies show a particularly low bias corrected replicate agreement within their top ranking sites. For strategies based on peak calling scores, this might be due to peaks corresponding to background binding regions. However, except for the *simple threshold* method, the top ranking sites of all other strategies show a lower agreement already before this bias correction in comparison to our method (Fig. S6a, S7a, S8a).

## PureCLIP captures strongest interaction footprints, not top ranking peaks

All previous crosslink site detection strategies, and in particular those based on peak calling scores such as 'Piranha^*sc*^ + CITS' and 'CLIPper^*sc*^ + CITS', call more sites in regions of high fragment density than PureCLIP in both basic mode and with the addition of co-variates (Fig. S6f and S7f). Further, the results show that these strategies also call far more sites within known common background binding regions than PureCLIP, even when not incorporating covariates. At the same time other strategies have far less 'bias corrected' agreeing calls between the two eCLIP replicates (Fig. 5). This indicates that the sites within the highest peaks are not necessarily corresponding to reproducible target-specific crosslink sites. These findings are in line with the results of the evaluation based on *bona fide* binding regions (Fig. 3), where strategies based on peak-calling scores ('Piranha^*sc*^ + CITS', 'CLIPper^*sc*^ + CITS') perform worse than corresponding strategies based on crosslink site detection scores ('Piranha + CITS^*sc*^', 'CLIPper + CITS^*sc*^'). In other words, most of the CITS sites within top ranking peaks are not located within regions matching the known binding characteristics of the proteins, and are thus likely to be false positives.

## Discussion

The detection of target-specific protein-RNA interaction sites from single-nucleotide resolution CLIP-seq data is a remaining challenge. Previous methods for the analysis of such data typically suffer from a large fraction of false positives, as they are sensitive to different sources of biases. Peak callers such as Piranha, which call regions enriched in read coverage without explicitly modelling read start counts at truncation sites, are prone to capture high background signal which does not originate from target-specific crosslink events. On the other hand, CITS calls sites with a significant fraction of read starts, but it can not distinguish, whether such sites are caused by target-specific crosslinks or by non-specific crosslinks within highly abundant regions. In addition, CITS does not account for biases, such as different transcript abundances or the crosslinking sequence bias, which can increase the number of false positives.

To overcome these limitations, we propose a new statistical approach called PureCLIP. PureCLIP calls crosslink sites considering both regions enriched in protein-bound fragments and the specifics of iCLIP/ eCLIP truncation patterns. It also explicitly models possible sources of bias, such as non-specific background signal and crosslinking sequence bias in order to reduce the number of false positives. Both these points, and in particular the incorporation of CL-motifs, which represent the non-specific crosslink sequence bias, are an innovation in comparison to existing methods.

A comprehensive evaluation based on simulated and real data has shown that, already in basic mode PureCLIP reaches a higher precision compared to previous strategies in almost all cases. Moreover, on real data the incorporation of input signals and CL-motifs additionally improves the precision of PureCLIP in capturing crosslink sites within *bona fide* binding regions. For the analysis of PUM2 and RBFOX2 data, it is worth noting that for the 50 top ranking sites (Fig. 3a,b, right panel) the precision of PureCLIP including the input signal is much higher in comparison to PureCLIP in basic mode or to previous strategies. The results indicate that the top ranking sites called by other strategies are likely to be caused by non-specific background signal, which is resolved by PureCLIP when incorporating input signal.

**Figure 5:**
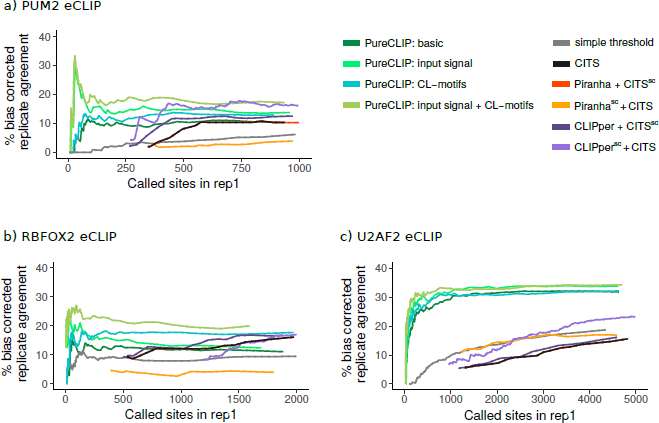
Agreement of called sites between replicates. On each eCLIP dataset (a: PUM2, b: RBFOX2, c: U2AF2), we report for each given number of called sites *x* (corresponding to a certain sensitivity threshold) in replicate 1, the percentage that were also called within the top *x* ranking sites in replicate 2 after correcting for different biases (see results, methods).

PureCLIP incorporating CL-motif scores strictly outperform all other strategies over all four datasets. In fact, PureCLIP's precision in this setting increases especially for the eCLIP datasets of proteins whose sequence motifs do not coincide with CL-motifs, namely PUM2 and RBFOX2 (Fig. 3). For RBFOX2 eCLIP data, the increase is particularly remarkable. This is also the only dataset where PureCLIP, without incorporating CL-motifs, shows a lower precision than strategies that make use of the CITS crosslink site detection score (Fig. 3b). The main reason is that PureCLIP in basic mode is more sensitive than CITS in also calling non-specific crosslink sites on this particular dataset (see Fig. S7d). In general a high sensitivity is desired, since we also want to detect crosslink sites for low-coverage regions, for example within lncRNAs or for proteins with lower binding affinity. In addition, false positives can be reduced by the incorporation of CL-motifs. Interestingly, when incorporating both input signal and CL-motifs simultaneously, PureCLIPs precision increases even further, highlighting the huge benefit of the incorporation of both covariates into the model.

Compared to previous strategies, PureCLIP achieves a higher agreement in calling RBP-bound sites between eCLIP replicates for *bona fide* crosslink sites. These are sites where the fragment coverage is enriched over the input signal, which do not overlap known background binding regions and, for PUM2 and RBFOX2, which are not located within CL-motif occurrences. Interestingly, the *simple threshold* method, which detects crosslink sites by applying a cutoff on the read start counts, has the worst performance of all on both simulated and real data, as expected, but by far the highest replicate agreement for all datasets when not explicitly accounting for biases. This indicates that, beside target-specific crosslink sites, other factors also contribute to this 'raw' replicate agreement, and that in order to obtain a meaningful evaluation of all methods, one needs to compute a 'bias-corrected' replicate agreement. These results also strongly suggest, that it would be valuable for the analysis of iCLIP/eCLIP data to explicitly include replicate information (as already suggested by [28]), but importantly that this needs to be done carefully while addressing possible sources of biases.

It is also important to stress that for all analysed eCLIP datasets PureCLIP calls far fewer crosslink sites within regions of high fragment density (Fig. S6f, S7f, S8f) and within known common background binding regions [21] (Fig. S6e, S7e, S8e) compared to all other strategies. This even holds for PureCLIP in basic mode. Taken together with PureCLIP's general higher precision, these findings demonstrate how important it is for the analysis of CLIP-seq data not only to call peaks, but also accurately model the counts of individual read starts, indicating potential truncation events. This unique feature of PureCLIP enables the distinction between target-specific interaction footprints and non-specific crosslink patterns within high abundant background binding regions.

Although the main objective of PureCLIP is to detect individual target-specific crosslink sites, it is sometimes desirable to identify larger binding regions for the protein under study. In the current version, PureCLIP can merge crosslink sites to larger binding regions based on their genomic distance. Further work will be needed in the future to address this task in a more sophisticated manner. However, results on simulated data demonstrate that individual crosslink sites also recover a large number of simulated binding regions with higher precision compared to the other strategies.

Currently, PureCLIP allows to incorporate covariates that influence either the emission probabilities of the pulled-down fragment density or the read start counts. Besides information on common background binding regions and replicate agreement, mappability information is a promising candidate that will be investigated further inside the PureCLIP model. Furthermore, the PureCLIP framework can be extended in the future to include covariates that directly influence transition probabilities between states, adapted to model other types of CLIP-seq diagnostic events or even other types of high-throughput data, such as RNA methylation.

## Conclusions

More and more high-resolution CLIP-seq datasets are being generated, but the precise determination of protein-RNA interaction sites from iCLIP/eCLIP has been challenging so far. Extensive evaluations demonstrated the superiority of PureCLIP over several previous strategies in detecting target-specific crosslink sites, both on simulated data as well as on real datasets. PureCLIP is able to precisely capture protein-RNA interaction footprints, while not relying on the highest peaks and being able to correct for biases, such as transcript abundances and crosslinking sequence preferences. It therefore provides a promising method to analyse these datasets, also for proteins with lower binding affinities or proteins binding to low abundant RNAs, such as lncRNAs.

## Materials and Methods

### Preprocessing of iCLIP/eCLIP datasets

We analysed four published eCLIP datasets targeting the proteins PUM2, RBFOX2 and U2AF2 and one iCLIP dataset targeting U2AF2 (see Additional file 1, Table S1 for details).

First, possible adapter contaminations at 3' ends were removed using TrimGalore on the iCLIP dataset [16] (v0.4.2, based on cutadapt), and by running cutadapt twice on the eCLIP datasets [18] (v1.12). The later was done in order to correct for possible double ligation events [30]. Subsequently, reads shorter than 18bp were discarded. Next, the reads were mapped against the human genome (hg19) using STAR [7] (v2.5.1b), a read aligner designed for RNA-seq data with setting '--alignEndsType End-ToEnd', '--scoreDelOpen −1' for gap penalty, and '--outFilterMultimapNmax 1' to discard reads mapping to multiple locations.

PCR duplicates were removed based on the read mapping positions and the random barcode sequences (also called UMIs). This is important, as PCR amplification rates are high, in particular for iCLIP datasets. To address this, we used UMI-tools [26], a network based de-duplicating method (in '--paired' setting), which is able to handle errors within barcode sequences.

All evaluated datasets come as two replicates. When assessing each method's capability to recover *bona fide* binding regions, we pooled the reads of the two replicates, whereas they were analysed separately when evaluating the agreement between called sites. Due to the differences in the two library preparation protocols [15] [30], we used either the 5'-end read (iCLIP), or the 3'-end read (eCLIP) of the sequenced fragment for the analysis.

## iCLIP/eCLIP-seq data simulation

In order the evaluate our method's performance, we developed a workflow to simulate realistic iCLIP-seq data starting from aligned RNA-seq data and known binding regions. The workflow simulates the main steps of the iCLIP/eCLIP protocols (see Fig. 6) as follows:

1. **Fragmentation:** To obtain RNA fragment lengths comparable to those of iCLIP experiments (30-300 bp, as described in [24]) we first simulate new fragment lengths using a normal distribution (mean: 165 bp, standard deviation: 50 bp).
2. **Binding regions:** We use genome-wide PUM2 motif occurrences computed with FIMO [11], in order to obtain a realistic distribution of binding regions (for details see Additional file 1, Section 2).
3. **Crosslink sites:** Within each binding region *i*, *c*_i_ crosslink sites are drawn uniformly (*c_i_ ∈* {1,…, 4}).
4. **Pull-down:** RNA fragments overlapping binding regions, are 'pulled down' with a certain rate. For this study we either used a pull-down rate of 1.0 or of 0.5, *i.e.* all or half of the overlapping fragments are used.
5. **Reverse transcription:** For each fragment, one of the following modifications can be applied to the 5'-end read:

a. The read start is shifted to one of the simulated crosslink sites within the current binding region according to a given truncation probability (set to 0.7).
b. The read start is shifted to any other position within the fragment according to a given offtarget truncation probability (set to 0.1).
6. **Size selection:** To obtain a broad range of cDNA lengths we keep reads with underlying fragment lengths between 30-140 nt (as recently recommended in [12]).

**Figure 6:**
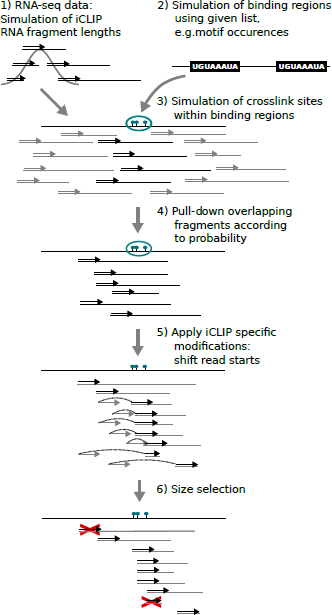
Simulation of iCLIP/eCLIP data.

In addition to the RBP binding signal, we also simulated background noise, that can be for example caused by sticky RNAs [9] or by the binding of non-specific background proteins [9]. We did this by applying the described steps on the list of known common background binding regions published in [21], while varying 'pull-down' rates, truncation probability and the number of crosslink sites within a region. We supplement those regions with reads randomly sampled from RNA-seq data (1%). Further details of the used simulation are described in Additional file 1, Section 2.

## PureCLIPs Hidden Markov Model

PureCLIP uses a Hidden Markov model (HMM) to infer crosslink sites from aligned single-nucleotide CLIP-seq data. At each position t, it utilizes two types of information (Fig. 7a): the pulled-down fragment density C_t_, which is used to infer whether the position is 'enriched' or 'non-enriched' in protein bound fragments, and the read start count K_t_, which is used to infer whether it is a 'crosslink' or 'non-crosslink' site. The four resulting hidden states are (1) 'non-enriched & non-crosslink', (2) 'non-enriched & crosslink' , (3)'enriched & non-crosslink' and (4) 'enriched & crosslink'. For the sake of clarity, we separate them into two state variables: one represents the enrichment state

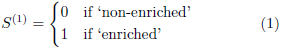

and one represents the crosslink state:

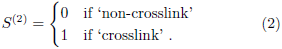

Our goal is to identify positions that are 'enriched & crosslinked' (see state (4) in Fig. 7a). While transitions between all four states are allowed, we model distinct emission probability distributions for each state.

### Joint emission probabilities and inference

We exploit the hierarchical structure of the two observed signals, i.e. the pulled-down fragment density (*C_t_*) and the count of read starts (*K_t_*), for specifying the model. First, we model the value of the fragment density *C_t_*, both for the 'non-enriched' and the 'enriched' state, with a left-truncated gamma distribution (LTG) use it for the fragment density values. We use a truncated gamma distribution as a practical solution as we do not want to consider positions which have a very low or no abundance.

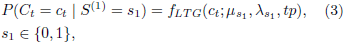

where *μ*_*s*1_ and λ_*s*1_ denote the mean and the shape parameter of the distribution and *tp* the truncation point. The parameters μ_*s1*_ and *λ*_s1_ need to be learned, while tp is fixed (see Additional file 1, Section 3.1 for details). The gamma distribution is a popular and flexible choice to model non-negative continuous values and thus we

**Figure 7:**
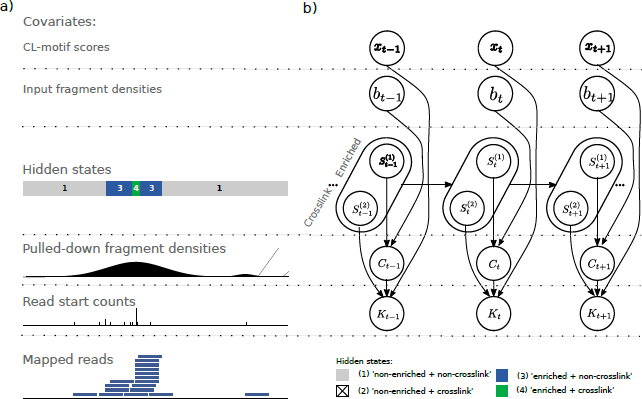
Summary of the HMM modelling framework. **a)** Starting from mapped reads (bottom), observations are deduced (read start count and pulled-down fragment density) and combined with additional covariates (top) to reconstruct the most likely sequence of hidden states (middle). **b)** Graphical representation of the corresponding non-homogeneous HMM.

When looking at the read start counts *K_t_*, we expect an increased count at 'crosslink' sites due to underlying truncation events. Therefore, we model the read start counts *K_t_* for both the 'non-crosslink' and the 'crosslink' state. For state *s*_2_ the probability to observe *k_t_* read starts is computed given the number of fragments *n_t_* covering the current region and the probability *p*_*s*2_ for each read to start at position *t*. *n_t_* is unknown but we can use a surrogate value *ň_t_* directly deduced from the positions pulled-down fragment density *c_t_* by a simple rescaling (for details see Additional file 1, Section 3.2.1). We model the emission probability distribution with a zero-truncated binomial distribution (ZTB)

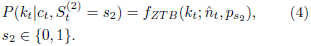

The probability parameters *p*_o_ and *p*_1_ need to be learned, where *p*_1_ reflects a protein specific truncation rate at 'crosslink' sites. A zero-truncated binomial distribution is preferred here as we do not want to fit the distributions to the large number of sites with no read starting.

Given the emission probability distributions for the four states, we compute the probability of a joint observation. Note that *C_t_* and *K_t_* are not conditionally independent, but since *n_t_* is directly deduced from *c_t_* the emission probability for the joint observation can be factorized accordingly (see Fig 7b for a graphical summary):

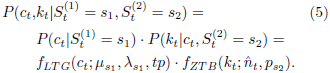

Finally, we use posterior decoding to determine the most likely hidden state for each position, and with that all 'enriched & crosslink' sites (*s*_1_ = *s*_2_ = 1). Each such called crosslink site comes with an associated score, namely the log posterior probability ratio of the first and second most likely state:

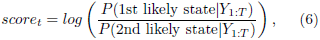

where *Y*_1:*T*_ denotes the observed data for all positions. In a second step, the called crosslink sites can be further combined to binding regions based on their distance.

We use the Baum-Welch algorithm [20] to learn the parameters of the HMM, i.e. the transition probabilities and the parameters of the four emission probability distributions (see Additional file 1, Section 3 for details on the implementation). For the parameter estimation of the emission probability distributions, only sites with at least one read starting are considered. Moreover, to reduce the computational costs, we trained the HMM on a subset of the chromosomes (Chr1 - Chr3 for pooled data, Chr1 - Chr6 for individual replicates). This had no impact on the quality of the estimates.

### Estimation of the pulled-down fragment density

To model the fragment density, we cannot use positions-wise read counts, since they will be strongly influenced by truncation events in the neighbourhood. Instead we apply a smoothing on the read start counts *c'* to estimate the density of pulled-down fragments at each position. This is done using a kernel density estimation (KDE) [19] with a Gaussian kernel function *K*. The latter assigns a higher weight to nearby read starts, while still considering read starts which are further away, thereby providing a better estimate for the underlying pulled-down fragment density. We compute the smoothed signal at position *t* using

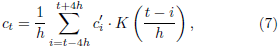

where *h* is the kernel bandwidth and positions within four bandwidths are considered.

## PureCLIPs non-homogeneous hidden Markov model

We incorporate position-wise external data as covariates into the HMM by using generalized linear models. Numerical optimization techniques are then used within the Baum-Welch algorithm to find the parameters that maximize the conditional expectation of the data.

### Incorporation of non-specific background signal

Without additional information, we assume that the fragment density *c*_t_ follows, for each enrichment state *s*_1_, a left-truncated gamma distribution

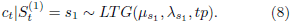

If an non-specific background signal is given, e.g. from an input control experiment, PureCLIP incorporates this as position-wise covariates into the model. This is done using a (left-truncated) gamma generalized linear model (GLM). The objective is to learn the correlation between the covariate *b* and the mean parameter *μ_s1_* of each enrichment state *s*_1_. The underlying multiplicative effect of the background signal *b_t_* on the expected mean *μ_s1,t_* is modeled using a log link function:

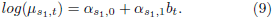

Note that for each enrichment state *s*_1_ we assume to have a constant shape λ_*s*1_ across the entire range of covariate values.

A numerical optimization is performed within the Baum-Welch algorithm to learn the parameters *α*_*s*1,0_, *α*_*s*1,1_ and λ_*s*1_ (see Additional file 1, Section 4.1). In this study we used the log fragment density of the input experiments as covariates, computed using a KDE with the same bandwidth as used for target fragment density, i.e. 50 bp.

### Incorporation of CL-motif scores

Without additional given information, the read start counts *k_t_* are modeled using a zero-truncated binomial distribution

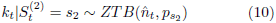

for each enrichment state *s*_2_. If we assume that we have learned *m* enriched CL-motifs from the input data (described in the next section), then we can compute for each position *t* and motif *i* ∈ 1,…, *m* a corresponding motif match score *x*_i,*t*_ >= 0, containing information about the positions crosslinking affinity. PureCLIP uses a logistic regression for each crosslinking state *s*_2_ to model the expected binomial probability parameter *p*_*s*2_ based on the position-wise CL-motif score *x*_i,*t*_

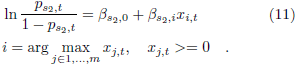

Since the majority of positions has no CL-motif match, i.e. a CL-motif score of 0, we compute *β*_*s*2,0_ using these sites as it was done in the basic PureCLIP model. Further, since we assume that each site only matches one CL-motif (i.e. the motif with the highest score is chosen) the parameters *β*_*s*2,1_,…, *β*_*s*2,*m*_ are learned independently of each other using the Brent's method (see Additional file 1, Section 4.2).

### Computation of CL-motif scores

The computation of position-wise CL-motif scores, that can be used as covariates by PureCLIP, is done in a preprocessing step. (1) We call crosslink sites on the input data using the basic version of PureCLIP (2) We run DREME (meme suite v4.11.3) [1] with the parameters '-norc -k 6 −4' on 10 bp windows spanning the called input crosslink sites, while using 10 bp windows 20 bp upstream and downstream as control (DREME uses Fisher's Exact test). (3) We use FIMO (meme suite v4.11.3) [11] with the parameters (--thresh 0.01 --norc) to compute occurrences of those motifs within the genome and their corresponding scores. If one position has overlaps with multiple CL-motif occurrences, the one with the highest score is chosen.

## Availability and implementation

PureCLIP is freely available as a command-line tool implemented in C++ using SeqAn [8], the GSL [10] and Boost [22]. OpenMP [5] is used for parallelization. PureCLIP is licensed under the GPLv3 and can be downloaded at **https://github.com/skrakau/PureCLIP**. Notably, as PureCLIP includes information from the transcriptomic neighbourhood, it is important to use a suitable reference sequence when mapping the reads. Thus, when analysing iCLIP/eCLIP data from proteins known to bind near exon-exon junctions on mRNAs, reads should be mapped directly against transcripts (e.g. as done in [12]). For the use of PureCLIP in conjunction with CL-motifs, a precompiled set of common CL-motifs (learned on pooled PUM2, RBFOX2 and U2AF2 input crosslink sites) is provided on the website.

Furthermore, we provide the framework to simulate truncation based CLIP-seq data, which requires mapped RNA-seq data and a list of binding regions. Additionally, background binding can be simulated based on previously published regions. The output of this simulation is a BAM file, containing both target and background reads, as well as BED files containing simulated binding regions and crosslink sites. The simulation workflow is also freely available under the GPLv3 license and can be downloaded at **https://github.com/skrakau/sim_iCLIP**.

## Comparison against previous crosslink site detection strategies

We compared PureCLIP against the following existing methods: *simple threshold,* CITS [31], Piranha [29] and CLIPper [17]. *Simple threshold* and CITS are methods to detect crosslink sites at single-nucleotide resolution and therefore can be directly compared with PureCLIP.

Piranha and CLIPper are strand-specific peak-calling methods and can not be directly compared to PureCLIP, therefore their performance was assessed in combination with CITS. In detail, we take the intersection between Piranha (p-value threshold 0.001) or CLIPper (default threshold) reported peaks with CITS crosslink sites (default p-value threshold) and score the resulting sites in two different ways: either according to the peak caller (referred to with the term Piranha^*sc*^ or CLIPper^*sc*^), or to CITS (referred with CITS^*sc*^). The assigned scores were used to assess the performance of the strategy for different sensitivity thresholds. Further details about the method's application and the parameter choice are described in Additional file 1, Section 5.

## Evaluation on real data based on *bona fide* binding regions

To assess the performance of the different strategies in detecting target-specific crosslink sites on the PUM2 and RBFOX2 datasets, we made use of the sequence motifs that were described for each of those proteins. FIMO [11] (--thresh 0.001 --norc) was used to compute genome-wide motif occurrences. Next, for each called crosslink site the distance to the closest motif start site was identified. The precision was defined as the percentage of all called sites within 2 bp of a motif occurrence (Fig. 3).

For the protein U2AF2 its known predominant binding site, which is ~ 11 nt upstream of 3' splice sites, was used for evaluation. Ensembl release 75 annotations were used to compute the distance of each called crosslink site to the closest 3' splice site. The precision is then defined as the percentage of all called sites that are located 11 ± 4 nt upstream of a 3' splice site.

## Computation of bias corrected replicate agreement

For the evaluation based on the replicate agreement, only sites with calls at the exact same nucleotide position in both replicates were considered as agreeing. In all comparisons, the replicate dataset with the larger library size was chosen as a reference for evaluation and will be referred to as replicate 2 in the following. We report for each given number of called sites *x* (corresponding to a certain sensitivity threshold) in replicate 1, the percentage that were also called within the top *x* ranking sites in replicate 2.

To compute the bias corrected replicate agreement, we count only sites that additionally (1) have sufficient enrichment over the signal obtained on input data, and (2) are not contained in common background regions [21] or in CL-motifs (for PUM2 and RBFOX2).

To determine the sites whose pulled-down fragment densities are enriched over the input, we chose an individual threshold for each protein dataset based on its distribution of log-fold enrichments (for details see Additional file 1, Section 7). CL-motif occurrences were obtained with FIMO as described previously, while using all matches with a score *>* 0. Common background binding regions were taken from [21], using only regions observed in at least 6 different CLIP-seq datasets, and extending them upstream and downstream by 200 bp.

## Competing interests

The authors declare that they have no competing interests.

## Author's contributions

SK and AM had the initial idea of the project. SK designed and implemented the PureCLIP model and performed all experiments. HR contributed to the design of the model. AM and HR supervised the study. SK, HR and AM wrote the manuscript. All authors read and approved the manuscript.

## Acknowledgements

We thank the ENCODE project consortium for making their data available and the authors of CLIPper for making their tool available on Github. We also kindly acknowledge Leon Kuchenbecker, Brian Caffrey and Stefan Budach for proofreading of the manuscript.

## Additional Files

### Additional file 1 — Supplementary Material

The Supplementary Material is provided as a .pdf file and it contains additional figures and more detailed information about the computational methods and the results.

